# The antibody repertoire of colorectal cancer

**DOI:** 10.1101/176768

**Authors:** Seong Won Cha, Stefano Bonissone, Seungjin Na, Pavel A. Pevzner, Vineet Bafna

**Author notes:** TCGA: The Cancer Genome Atlas Project CPTAC: Clinical Proteomic Tumor Analysis Consortium TIL: tumor infiltrating lymphocyte Ig: immunoglobulin SdB: ‘split’ de Bruijn dB: de Bruijn FR: framework region CDR: complementarity determining region IMGT: the international ImMunoGeneTics information system COAD: colon adenocarcinoma DP: Digital Proteomics SAAV: Single Amino-Acid Variant.

## Abstract

Immunotherapy is becoming increasingly important in the fight against cancers, utilizing and manipulating the body’s immune response to treat tumors. Understanding the immune repertoire – the collection of immunological proteins – of treated and untreated cells is possible at the genomic, but technically difficult at the protein level. Standard protein databases do not include the highly divergent sequences of somatic rearranged immunoglobulin genes, and may lead to missed identifications in a mass spectrometry search. We introduce a novel proteogenomic approach, AbScan, to identify these highly variable antibody peptides, by developing a customized antibody database construction method using RNA-seq reads aligned to immunoglobulin (Ig) genes.

AbScan starts by filtering transcript (RNA-seq) reads that match the template for Ig genes. The retained reads are used to construct a repertoire graph using the ‘split’ de Bruijn graph: a graph structure that improves upon the standard de Bruijn graph to capture the high diversity of Ig genes in a compact manner. AbScan corrects for sequencing errors, and converts the graph to a format suitable for searching with MS/MS search tools. We used AbScan to create an antibody database from 90 RNA-seq colorectal tumor samples. Next, we used proteogenomics analysis to search MS/MS spectra of matched colorectal samples from the Clinical Proteomic Tumor Analysis Consortium (CPTAC) against the AbScan generated database. AbScan identified 1, 940 distinct antibody peptides. Correlating with previously identified Single Amino-Acid Variants (SAAVs) in the tumor samples, we identified 163 pairs (antibody peptide, SAAV) with significant co-occurrence pattern in the 90 samples. The presence of co-expressed antibody and mutated peptides was correlated with survival time of the individuals. Our results suggest that AbScan(https://github.com/csw407/AbScan.git) is an effective tool for a proteomic exploration of the immune response in cancers.

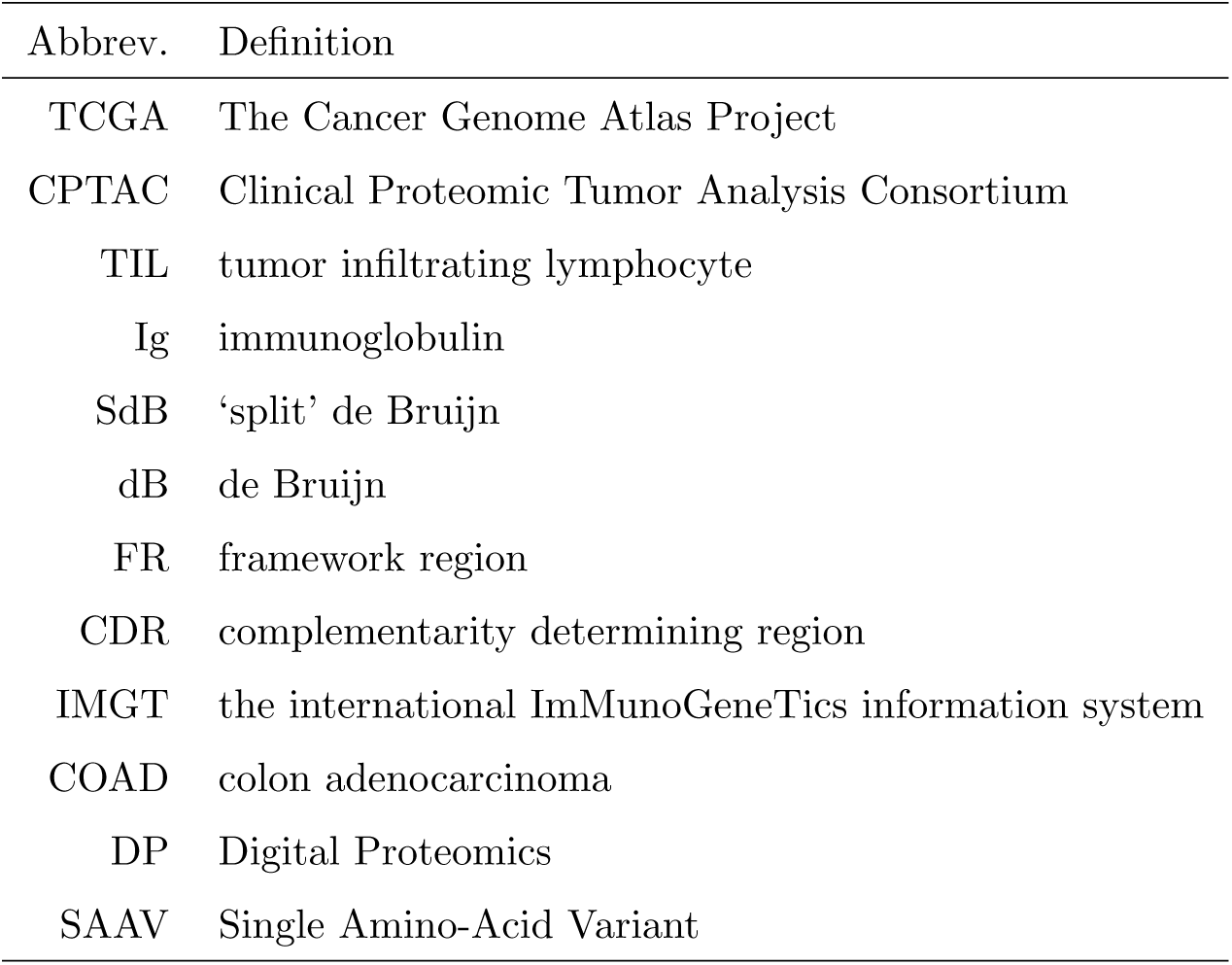

## 4 Introduction

Cancer immunotherapy, which attempts to tackle cancer using the body’s own immune response, has been very successful in boosting the survival rates of patients with leukemia and other blood cancers [1–3]. This field of research is expanding rapidly, and has been extended to include other cancer sub-types, including solid tumors [4–6].

Immunotherapy is more specific than generic typical cancer treatments targeting fast-growing cells directly. It can take the form of cancer vaccines (neoantigens that stimulate an immune response) [7, 8], monoclonal antibodies, which target cancer cells expressing specific (neoantigenic) proteins [9] or immune checkpoint inhibitors that activate suppressed immune cells [10–12]. The development of new forms of cancer immunotherapy could be greatly helped by knowledge of the cancer specific immune response, especially in understanding the antibodies and neoantigens specific to cancer.

This is a challenge because of the millions of distinct antibodies that are circulating in the blood. We still have only limited knowledge of the antibody responses that target individual disease-related antigens and epitopes. There are only a few known examples in infectious disease [13] and autoimmune disease [14]. On top of that, recent methods that characterize the antibody repertoire use serum or plasma samples as their source for antibody analysis. However, the antibodies in these samples include the pool of all antibodies binding to multiple antigens, as well as the antibodies produced by numerous previous immune responses [15–19]. Screening the antibodies based on their binding to pre-selected antigens may also not work, as all possible neoantigens existing in a sample cannot be known, and some important antigens may be post-translationally modified [20, 21] or cleaved [22].

Another approach to understanding the antibody repertoire is by isolating the B-cells that respond to a target immunogenic antigen. Plasmablasts [23, 20], memory B cells [24–26], and tissue infiltrating B cells [27–29] have been used to characterize the functional antibody repertoire [30, 31]. The method works, but it requires a dedicated workflow to isolate the B-cells and sequence the antibody clones. Here, we propose a more direct method for discovering antibody peptides in tumor samples.

Recently, we and others have developed pipelines for identifying mutated peptides expressed specifically in cancer [32–34]. In our approach, we mine a general transcript resource (such as The Cancer Genome Atlas Project) to extract transcript sequences, identify novel mutations, and junctions, then encode them into a complex database. This database is then searched via a proteogenomics approach, to identify peptides that are seen only in tumor proteome samples. Interestingly, our initial search of the CPTAC colorectal tumor samples identified a number of antibody peptide sequences [32]. Supplemental Fig. S1 shows the example of some antibody peptides identified in the search. At the time, there were questions regarding the provenance of the discovery, as we did not expect to find antibody peptides in colon tissue. They could be antibodies from tumor infiltrating lymphocytes (TIL), circulating antibodies from blood contamination, encoding general proteome variation, or even mis-identifications. Moreover, our databases were not specifically designed to capture Ig regions, so we were only identifying peptides from some of the annotated Ig genes on the human reference.

AbScan is a new tool for identifying all antibody (Ig) peptides in a sample by searching mass spectral datasets against RNA-seq datasets. AbScan is a proteogenomic tool that scans transcript and genomic data, preferably, but not exclusively from the same samples as the proteomic data; it creates specialized antibody sequence databases that can search tandem mass spectra. As the antibody sequences are hypervariable, identifying and characterizing transcripts encoding Ig genes is a challenging endeavor. We devised a special construct called the ‘split’ de Bruijn (SdB) graph to encode all Ig transcripts in a compact fashion, then show the power of this approach compare to other methods. AbScan also uses a customized pipeline to search these antibody databases and identify expressed antibody peptides, while controlling for false discoveries. We evaluated sensitivity and specificity of AbScan by benchmarking it on simulated datasets, pure antibody mixtures, normal colon tissues, and colorectal cell-lines. We further applied AbScan to 90 colorectal samples from the CPTAC project and demonstrated that the antibody repertoire was characterized by significant co-occurrence pattern in 163 pairs of antibody peptides and Single Amino-Acid Variant (SAAV) pairs, and the co-occurring pairs were correlated with patient survival.

## 5 Experimental Procedures

### Experimental MS data sets, sequence databases, and search parameters

We analyzed four spectral datasets, which have been described in previous work.

- 90 colorectal tumor samples [35]
- 30 normal colon biopsies [35]
- Colon cancer cell-lines LIM1215, LIM1899, and LIM2405 [36], and
- Purified polyclonal antibody mixture [37].

We searched each tandem mass spectra against three different databases. These included:

- Ensembl database version GRCh38 [38]
- The ‘split’ de Bruijn (SdB) graph based database driven by the method described in this paper, and
- A de Bruijn (dB) graph based database [34].

We used MS-GF+ (version 1.1.0) [39] with the following parameters: parent mass tolerance of 20 ppm, and allowed post-translational modifications of fixed carbamidomethyl C and optional oxidized Methionine. Common contaminants were excluded. ProteoWizard (v3.0.3827) [40], and ReAdW (v1.1 and v4.3.1) [41] were used for the peaklist-generating software. Number of missed and non-specific cleavages permitted was 1.Trypsin was used to generate peptides for three colorectal data sets, and Trypsin, Asp-N, Chymotrypsin, Elastase were used for the purified polyclonal antibody mixture data set. A multi-stage FDR (See Experimental Procedures–‘Multi-stage search’) was applied to identify the PSMs from the SdB and dB driven databases (See Supplemental Table 1).

**Table 1:**
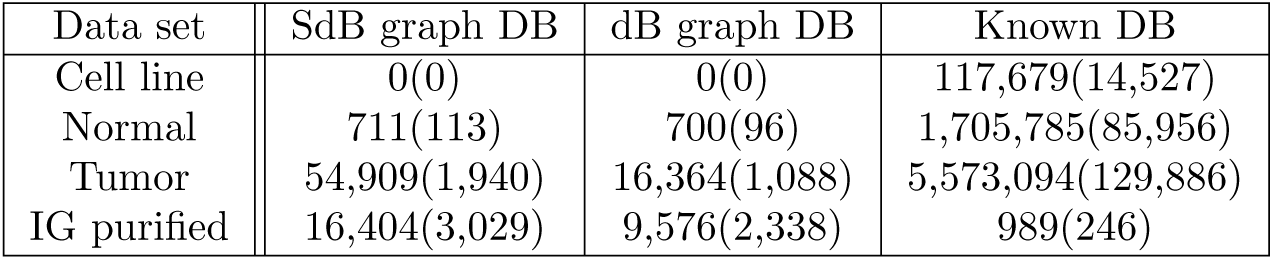
Number of identified PSM(peptides)

| Data set | SdB graph DB | dB graph DB | Known DB |
| --- | --- | --- | --- |
| Cell line | 0(0) | 0(0) | 117,679(14,527) |
| Normal | 711(113) | 700(96) | 1,705,785(85,956) |
| Tumor | 54,909(1,940) | 16,364(1,088) | 5,573,094(129,886) |
| IG purified | 16,404(3,029) | 9,576(2,338) | 989(246) |

### Database construction using ‘split’ de Bruijn (SdB) and de Bruijn (dB) graphs

AbScan constructs the ‘split’ de Bruijn (SdB) graphs for multiple RNA-seq datasets from tumors. A de Bruijn (dB) is constructed for a fair comparison. Followings are the steps to generate a custom MS searchable database:

1. **Read filtering**. Filter out all RNA-seq reads not sampling an Ig gene.
2. **SdB graph construction**. Create a SdB graph based database from filtered reads.
3. **Error correction**. Identify and eliminate sequencing errors.
4. **FASTA database construction**. The SdB graph is used to generate an MS searchable FASTA formatted database, as well as scripts to identify the context of the peptide on the antibody sequence.

For comparing the performance of SdB graphs to dB graphs, we used an implementation of dB graphs customized for the discovery of antibody peptides [34].

### Read filter

All antibodies are a combination of relatively fixed framework (FR), and hyper-variable complementarity determining regions (CDR), with the order given by ‘FR1, CDR1, FR2, CDR2, FR3, CDR3, FR4’. The typical lengths of CDRs in human are 15 to 30 nt for CDR1 and CDR2, and 24 to 36 nt for CDR3 [50]. On the other hand, lengths of RNA-seq reads in our datasets varied from 76 to 100 nt. Therefore, we expect most RNA-seq reads to cover some part of a framework region, and could use this to filter RNA-seq reads from Ig genes. In addition, we employed keyword matching to recover non-mapped Ig gene encoding reads by creating a list of *k*-mer sequences from all Ig genes in the IMGT reference [42], and selecting all reads that matched one of the *k*-mers.

An appropriate value of parameter *k* was determined by comparison with decoy data obtained by reversing the IMGT reference sequences. As *k* is made smaller, we can quantify the false matches by the number of reads that match decoy *k*-mers. For any value of *k*, the false discovery rate is given by

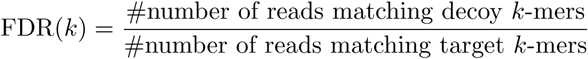

We selected the smallest value of *k* that resulted in a FDR below 1%, *k* = 19 was used for filtering (Supplemental Fig. S2). Quality filtering was applied additionally to trim the part of the poor quality reads. We trimmed the 3′ end of the reads if their quality threshold were less than the threshold value (10). We excluded the read if the trimmed part was longer than 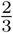 of the read length or the overall quality of the reads were below than the threshold value (25).

### SdB graph construction

Typical de Bruijn (dB) graph construction is as follows: Given a set of reads, the de Bruijn (dB) graph for this set is defined as follows: each *k*-mer from reads is a node in the graph. Nodes *u* and *v* are connected by an edge, if there exists a (*k* + 1)-mer in reads whose *k*-suffix is *u* and whose *k*-prefix is *v*. dB graphs are a powerful construct because they help remove redundancy in read coverage, and can be efficiently constructed without the need to compare all pairs of reads to test for overlap [43–45]. In the ideal case, each of the Ig genes is a path in the graph, and each path in the graph corresponds to a putative Ig gene. Errors can arise in dB graph construction if two unrelated reads share the same *k*-mer (we denote these as ‘false-positive’ overlaps), or if reads from the same molecule do not share a *k*-mer due to sequencing errors (false-negatives).

False edges in the dB graph can also arise due to repeated *k*-mers. Specifically, a repeated substring of size greater than or equal to *k* will lead to false edges in a *k*-mer based dB graph. The error could be controlled by selecting larger values of *k*, but that would result in a higher false-negative rate. We reasoned that the exact match requirement in *k*-mer dB graph is restrictive. For example, consider two reads that overlap over 40 bp. The probability that this overlap contain *k* = 30 consecutive nucleotides with no error in both reads is 65.5%. On the other hand, the probability that this overlap contain a 30 consecutive nucleotides with at most one error is 93.0%. Therefore allowing for an approximate match improves sensitivity from 65.5% to 93.0%. See Supplemental Method – ‘Analytical comparison of SdB and dB graphs’ for a rigorous analysis.

An alternative aproach is to do an error-correction prior to matching. BayesHammer uses a Bayesian approach on 1-neighborhoods of *k*-mers to correct reads, before constructing a *k*-mer dB graph [46]. In Ig genes, however, we use RNA data to identify variation, and the variable coverage makes it difficult to distinguish sequencing errors from true genetic variation. The SdB graph handles this problem of non-uniform coverage through correction on local nodes similar to the IGdb, Trinity and IDBA-tran [47, 48, 34].

On top of that, SdB graph applies a binning technique to solve the approximate matching problem efficiently. To obtain 1-neighborhoods of *k*-mers, we need the pairwise distance of every existing *k*-mers observed from the reads, which may increase the computation time for large value of *k*. We divided the *k*mers into two parts: one (*r*-mer) for binning, and the other (*𝓁*-mer) for 1-neighborhood testing. The size of the bin decides the average number of nodes required the pairwise distance computation. Note that the SdB graph is a generalization of prior approaches with bin size of *k* (respectively, 0) corresponding to a standard dB graph (respectively, BayesHammer like graph). In our tests, we did some empirical tests to choose *r* and *𝓁* and found the performance to be robust to difference choices. Therefore, we worked with *r* = 10, *𝓁* = 20, leaving the optimization of parameters to future work. However, to allow for fair comparisons, we tested the SdB graphs against dB graphs using a range of values of *k*.

Given *r, 𝓁*, we build a SdB graph as follows:

1. Each node initially corresponds to a distinct (*r* + *𝓁*)-mer from the read. Node *u* = (*x, y*), where *x* is a length *r* of prefix of the node, and *y* is a length *𝓁* of suffix of the node.
2. Consider nodes *u* = (*x, y*), and *v* = (*x′, y′*). We connect *u* and *v* by an edge, if the *r* + *𝓁 -* 1 suffix of *u* matches the prefix of *v*, and a read matches the combined sequence. The weight of an edge (*u, v*) is the number of reads that contain the combined sequence. The weight of node *u* is the maximum of the sum of incoming or outgoing edge-weights. This operation mimics a standard dB graph construction.
3. Consider nodes in order of decreasing weights, and repeat the following until no node is left:

a. Pick node *u* = (*x, y*). For all nodes *u′* = (*x′, y′*), merge *u′* with *u* (and remove from further consideration) if *d*_*h*_(*x, x′*) = 0, *d*_*h*_(*y, y′*) *≤* 1 and *u* is the heaviest, where *d*_*h*_(*x, x′*) is hamming distance between *x* and *x′*. Merge any multi-edges into a single edge of weight equal to the sum of the weights of the merged edges. Note that the actual implementation speeds this computation by hashing on the prefix strings.

The construction of a SdB is illustrated in Fig. S3. A (3, 3) SdB graph successfully compacted the data with no false-positives or false-negatives except due to sequencing error. As Fig. S3 shows, there are two distinct paths, corresponding to the two genes. However, due to sequencing error, we see a small branching in gene 1. This can be controlled by an error correction procedure, described in the next section. In contrast, Supplemental Fig. S4 shows examples of dB graphs with the choice of *k* = 4 and *k* = 5 using same reads. A 4-mer dB graph connected false edges at node ‘*GAAT*’, producing false paths combining gene 1 and 2. On the other hand, a 5-mer dB graph failed to connect edges in both genes, and neither gene could be represented by a single path. In the Results section, we systematically compare the performance of SdB graphs and dB graphs.

### Error correction

Sequencing errors also result in false overlaps. An error towards the end of the read (within *k* nucleotides) leads to a ‘tip’ in the dB graph, while an error in the middle of the read leads to a ‘bulge.’ Subsequent to its construction, the SdB graph can be viewed as a regular (*r* + *𝓁*)-mer graph; graph simplification methods, such as tip clipping and bulge removing can be applied. For transcript assembly, uniform coverage pruning may delete some true sequences, so we use a proportional approach to rescue lower-abundance transcripts similar to the one used by IGdb, Trinity and IDBA-tran [47, 48, 34].

Assuming for simplicity that sequencing errors are independent and identically distributed with ∈_*s*_ denoting the nucleotide error probability. The number of reads matching a specific *k*-mer is proportional to (1 − *∈*_*s*_)^*k*^. On the other hand, the number of reads matching a *k*-mer with a mismatch at a specific position is proportional to 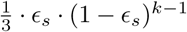. The expected ratio of read depths of the true edge to any false edge is given by

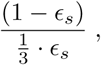

The expression is usually ≫ 1, for typical values of ∈_*s*_ *∼* 1%. Therefore, sequencing errors can be overcome as long as the sequence coverage is high enough.

AbScan differentiates true mutations from sequencing errors using the same idea. In ideal case, any genes conveying mutations are regarded as separate genes and the graph maintains separate paths. However, if two genes are separated only by a few polymorphisms, then the graph may merge some nodes in paths. For SdB graphs, an exact match requirement for the *r*-mer would result in a bulge where one collection of *r*-mers carry the mutation, and the other collection contains *r*-mers carrying the reference nucleotide. In the case of a true mutation, these bulge would be well-supported by reads, and not removed during error correction.

### FASTA conversion

To use the SdB graph to construct a FASTA database, we associated a sequence with each node. The sequence of the source is the *r*-mer; the sequence associated with the sink is the last nucleotide of its *r*-mer concatenated with its *𝓁*-mer. For all non-source, non-sink nodes, the associated sequence is simply the last nucleotide of the *r*-mer. The sequence of a path in the graph is the concatenation of sequences associated with nodes on the path. A compact FASTA database is constructed from the SdB graph by enumerating the paths as described. The sequences in the path were converted to the amino-acid FASTA format to generate a database for the MS/MS database search tools, using the SpliceDB tool [32] for this conversion. 69.3MB of FASTA form amino acid database was created, concentrating on the antibody sequence generated from 162.7GB of RNA-seq bam files.

### Multi-stage search

The antibody database adds some noise to the search and it is possible that a PSM to a known peptide has a better score against an antibody peptide, leading to false identification. As a conservative strategy to avoid false identifications, we use a modified multi-stage search [33]. We first searched all sepctra using MS-GF+ against a known protein database (Ensembl version GRCh38) [38]. All PSMs identified as a non-Ig known peptide from the spectrum level 1% FDR search of the Ensembl database were excluded from the second search. Spectra that could not be matched were searched using MS-GF+ against the antibody database, using a target-decoy strategy with 1% spectrum level FDR.

### Comparison to rnaSPAdes

As transcriptome assembly is a well established research area, we built database using a popular transcriptome assembly tool, rnaSPAdes[49] to compare with AbScan. To make a fair comparison, we applied the identical read set for assembly. SPAdes version 3.9.0 was used with options “– only-assembler” and “–rna”. The output nucleotide sequence translated to FASTA form amino-acid sequence for MS/MS search.

### Identifying Antibody peptide location

For all identified antibody PSMs, we found the most likely position in the antibody structure. To do this, we recovered the nucleotide sequence of the peptide from the SdB graph, then compared it to sequences with the IMGT sequence to find the best matched position of each PSM to IMGT reference sequences. Finally, we incorporated gaps to the position using IMGT multiple sequence alignments to get a normalized position. Fig. 1 shows the expected position of the peptides we identified from the colorectal tumor MS/MS data and polyclonal antibody MS/MS data. Each horizontal black line represents the distinct peptide sequence. Peptides that do not map to IMGT reference sequences are not displayed.

**Figure 1:**
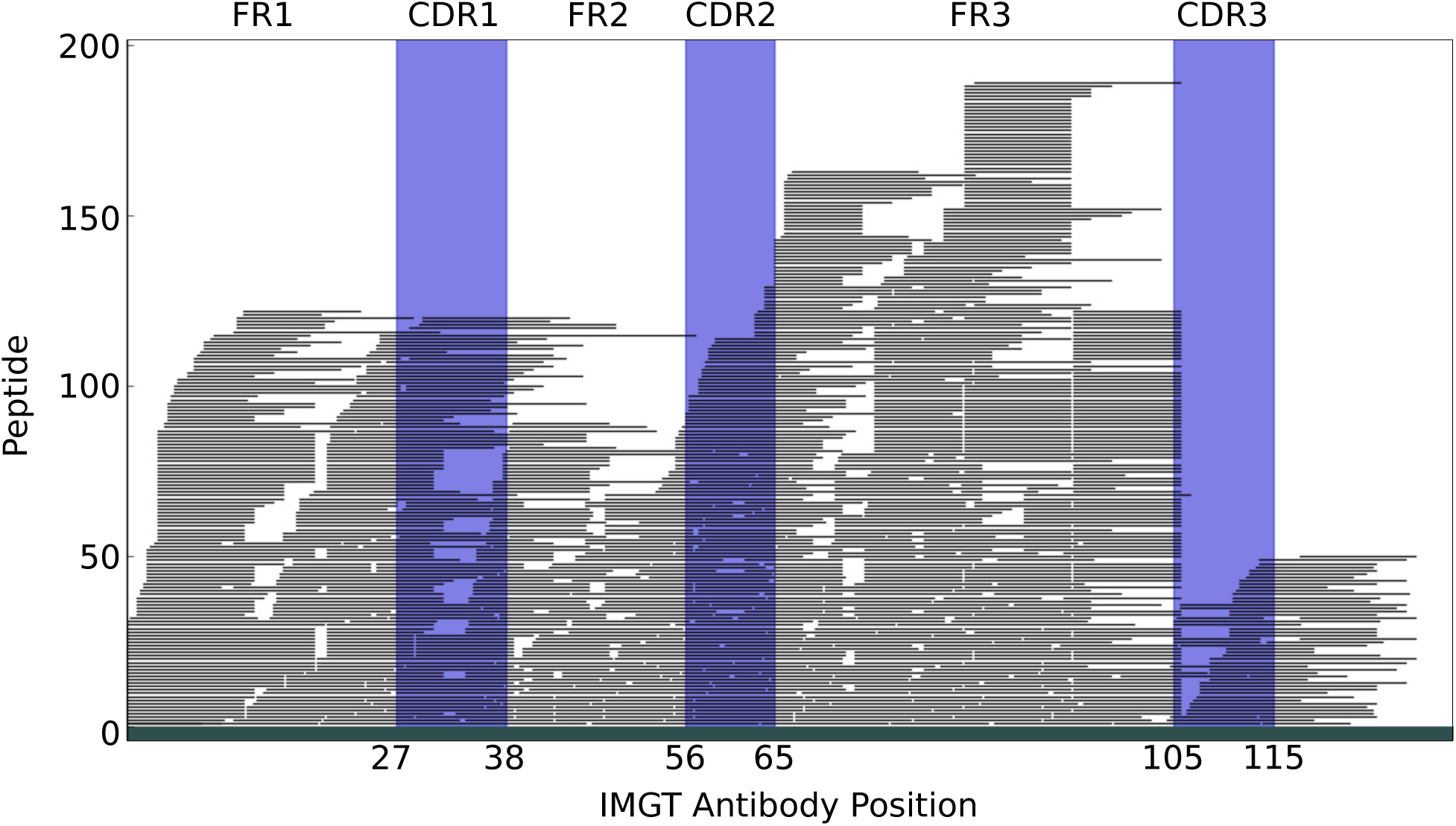
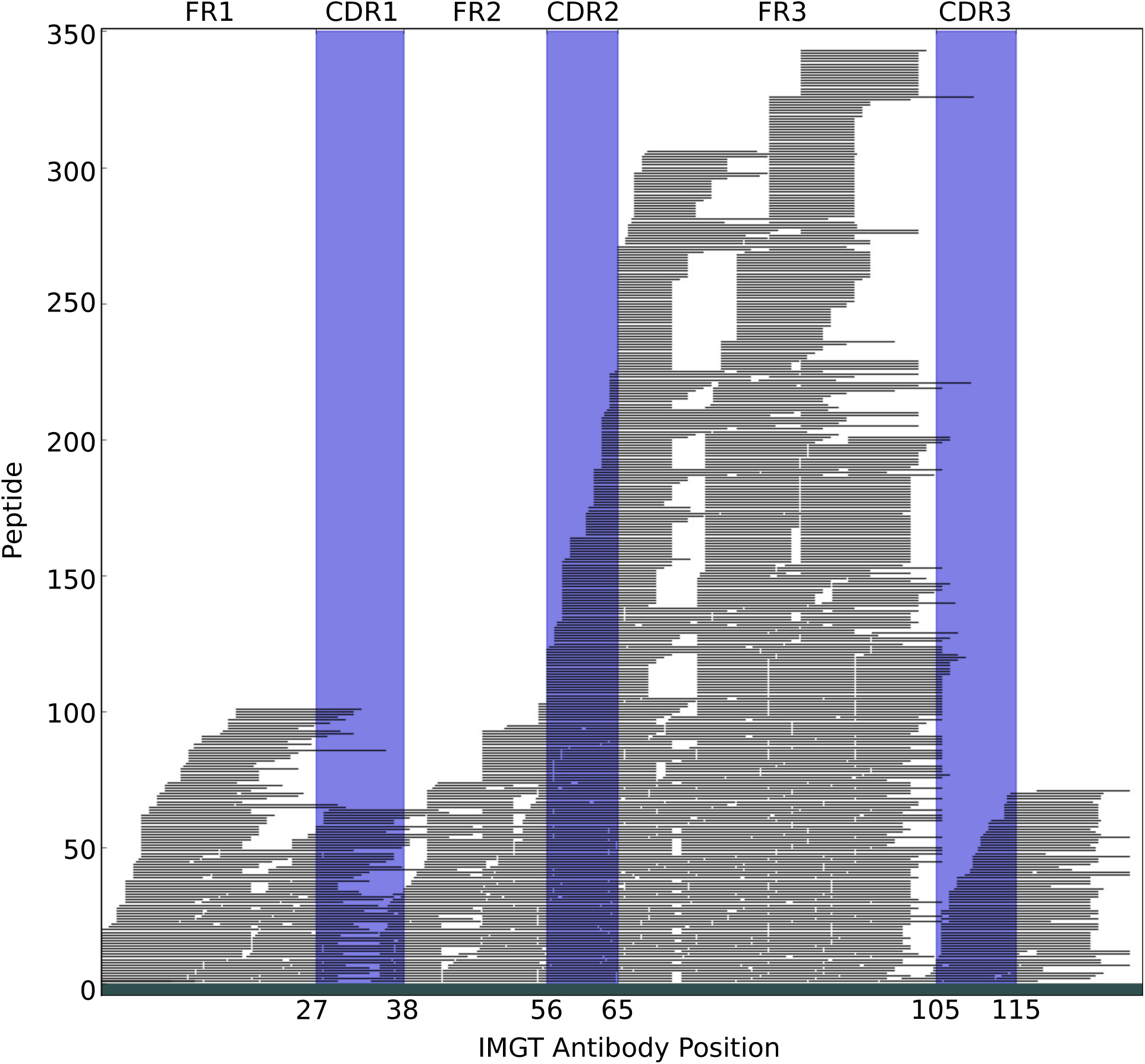
Relative locations of identified antibody peptides. Each horizontal black line represents a distinct peptide sequence. Trypsin was applied for the colorectal tumor MS/MS spectra assessment, and four different enzymes were applied for polyclonal antibody MS/MS spectra assessment. Both spectra sets were searched against the same antibody database constructed using tumor RNA-seq reads driven by TCGA. **(a)** Antibody PSMs from colorectal tumor MS/MS data. **(b)** Antibody PSMs from polyclonal antibody MS/MS data.

### Statistical test for antibody enrichment

The two stage search resulted in PSM identifications with spectra matching ‘known peptides’ and antibody peptides (SdB database). If in some sample, the MS/MS data was known to not contain any antibody peptide (e.g., cell-line), then any PSM in the SdB database corresponds to a false identification. The number of false identifications is expected to grow linearly with the number of known peptide identifications. Therefore, we considered the fraction

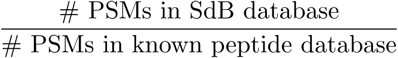

for all MS/MS data, and considered the Null hypothesis that this fraction was constant in all cases, colorectal tumor, colorectal normal, and colorectal cell-lines. To calculate the p-values, we applied Pearson’s χ^2^ test in a 2 × 2 contingency table.

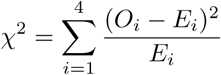

where

*χ*^*2*^ = Pearson’s cumulative test statistic,

*O*_*i*_ = Number of observation of type i

*E*_*i*_ = Theoretical frequency of type i

The *p*-value was calculated from the *χ*^*2*^ distribution table.

### Antibody and SAAV peptides correlation test

Consider a table, where columns correspond to samples (each column is a different sample), and rows correspond either to SAAV peptides (possible antigens) or to antibody peptides. The cells mark the presence or absence of the peptides in the specific sample. For any pair of antibody and SAAV peptides, we used the Fisher’s exact test to measure correlation of occurrence. As a large number of pairs were used, we used a target-decoy based approach to compute the false discovery rate for any *p*-value cut-off.

For each row, the columns were permuted independently so that any correlation between two rows (an antibody peptide, antigen pair) was just by chance, and a Fisher exact test was used to compute the correlation between all pairs. Highly correlated pairs of antibody and candidate antigen peptides were identified by applying a 5% FDR threshold.

### Measuring immune response

Measurement of the immune response for each individual was accomplished by counting the number of antibody PSMs. As the total number of spectra and their quality were not identical for every sample, the antibody PSMs were normalized by the total number of PSMs to the ‘known database’ search. Fig. 2(c) shows the distribution of the normalized immune response of each individual in both tumor and normal samples. We simply took the top 45% and bottom 45% group of individuals in terms of their normalized immune response.

**Figure 2:**
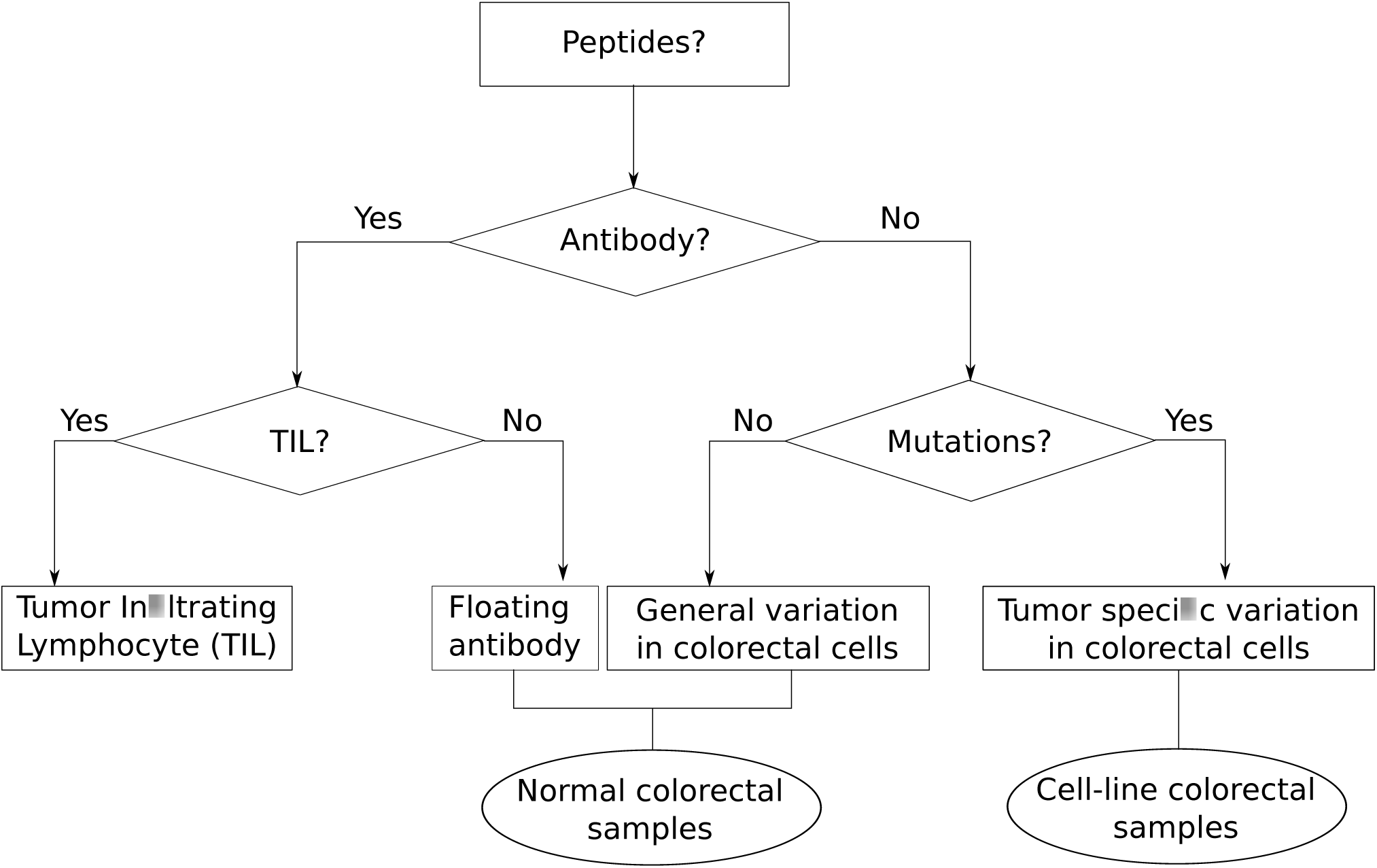
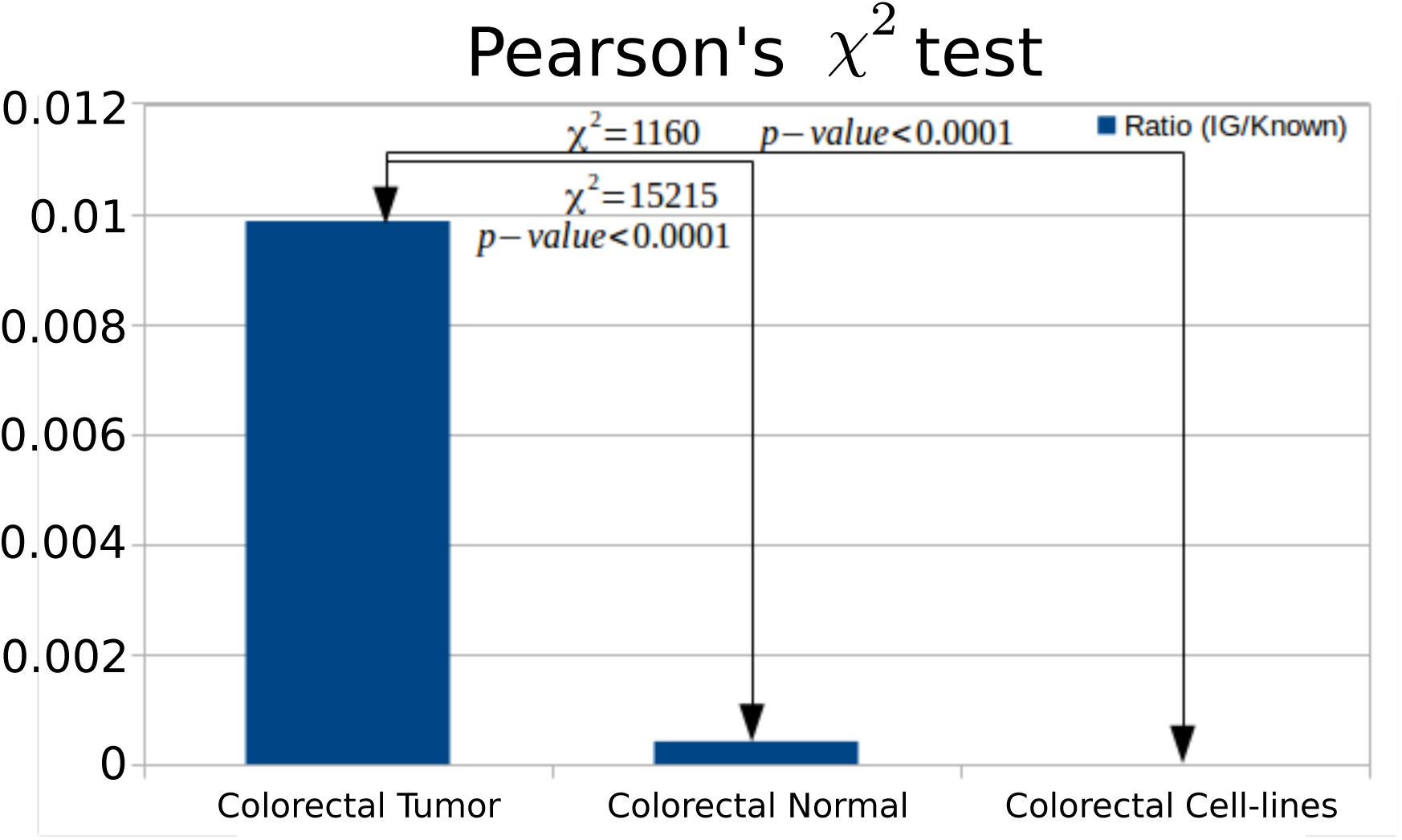
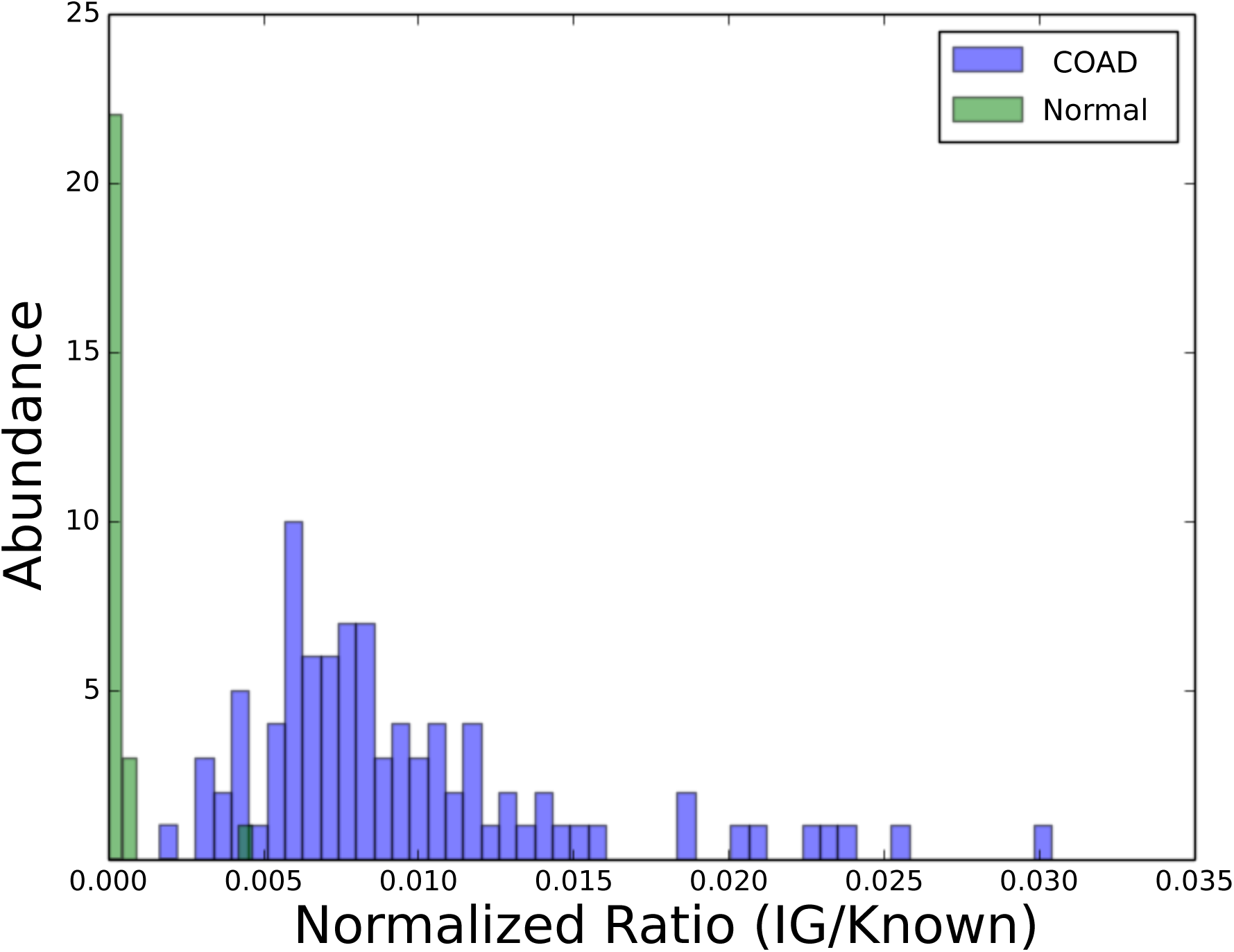
Comparison of identified antibody PSMs per experiment and sample. (a) The source of antibody peptides in different samples. PSMs that match non-reference peptides are either mutations or antibody peptides. Antibody peptides should not be observed in cell-lines. However, floating antibodies could be observed in normal colorectal samples. Antibodies from Tumor infiltrating lymphocytes should only be observed in tumor samples. (b) Occurrence of antibody peptides in tumor, normal, and tumor derived cell-lines are significantly different for MS/MS spectra of tumor, normal, and cell-line colorectal samples. Each spectra set were searched against the Ensembl GRCh38 protein database[38] and a custom antibody database. The number of PSMs identified as antibody peptides were 54*K* (*colorectal tumor*), 711 (*colorectal normal*), and 0 (*Cell-lines*). The PSM counts were normalized against the number of PSMs to known peptides. 5.5*M* in *colorectal tumor*, 1.7*M* in *colorectal normal*, and 0.1*M* in *Cell-lines*. The normalized ratios suggest that a significantly larger fraction of the colorectal tumor PSMs are antibody peptides, compared to the other two data-sets (Pearson’s *χ*^2^ *p*-val *<* 10^−4^). (c) The distribution of the number of samples carrying a normalized fraction of antibody peptides. COAD samples carry a higher fraction of antibody peptides.

### Survival rate comparison

We designed a method that takes a collection of peptides, and samples, groups samples based on co-occurrence of peptides, and tests if the individuals groups have different survival times. Specifically, we used the following strategy:

1. Represent each peptide *p* as a binary vector **p** over all samples with **p**_*i*_ = 1 (respectively, **p**_*i*_ = 0) indicating the presence (respectively, absence) of peptide *p* in sample *i*.
2. Cluster the peptide vectors into two groups (arbitraily labeled +, −) using 2-means clustering.
3. Assign score *S*_*i*_ to each sample *i* using:
*S*_*i*_ = (Number of ‘+’ assigned peptides in sample *i*) − (Number of ‘−’ assigned peptides in sample *i*)
4. Pick two sets of samples: Bottom (45%) of all samples with the lowest score, and top (45%) with the highest score.
5. Perform the Kaplan Meier log-rank test on the two groups of samples to test for correlation with clinical outcome.

We performed this test using all antibody and mutated peptides that significantly co-occurred in the samples exceeding a 5% FDR threshold of correlation test. The test statistic could include some unknown bias, and it wasn’t clear if they followed the *χ*^*2*^ distribution used to compute a *p*-value. To test this, we set two groups of patients where each group included 45% of random samples without replacement, and then we calculated the test statistics of two groups of random patients by log-rank test. We repeated the process 10, 000 times to create the distribution of test statistics (Supplemental Fig. S5).

## 6 Results

### Analytical comparison of SdB and dB graphs

We compared the performance of SdB graphs versus dB graphs using both analytical methods as well as empirical data from simulations. Let *p*_*s*_ denote the probability that a randomly chosen pair of nucleotides are identical. Thus, the probability of a false *k*-mer match is *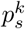*. To allow for fair comparisons, parameters *r, 𝓁, k* were selected so that the probability of an (*r, 𝓁*) match between unrelated reads in a SdB graph is the same as the probability of a *k*-mer match in dB graph. Specifically (Supplementary Methods–‘Analytical comparison of SdB and dB graphs’),

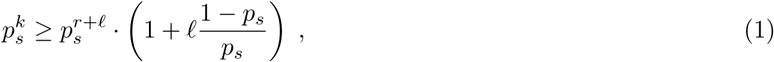

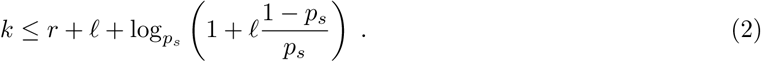

For any *r, 𝓁*, we chose *k* to be the largest value satisfying constraint 2. We also computed the probability of false overlaps. (See Supplemental Methods–‘Comparison between the SdB and dB graph mathematically’). Using these calculations, we can show that SdB graphs have significantly lower false negative rates compared to dB graphs. For example, let *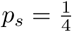*. When *r* = 10, *𝓁* = 20, the choice of *k* = 27 equated the false overlap rates for both methods at 5.55 · 10^−^17. However, for an overlap of 40 bp, *ε* = 0.01, we computed a false negative rate of 13.9% for the dB graphs versus a rate of 2.2% for the SdB graphs.

To test these theoretical results, data was simulated by generating the 100, 000 overlapping regions, of length from 30 to 100, with uniform sequencing error rate *ε*. Reads were connected by a path in the dB graph, if there was at least one *k*-mer consecutive sequence without an error. Similarly, they were connected by a path in a SdB graph, if there was an (*r* +*𝓁*)-mer in which the first *r* nucleotides had no error and the following *𝓁*-mer had at most one mismatch. Supplemental Fig. S6 showed a complete concordance between theoretical and simulated results. The sensitivity for all methods increases with length of overlap and decreases with higher *ε*. SdB graphs consistently outperform dB graphs.

### Comparison on simulated Antibody reads

To provide a more direct comparison of the performance of SdB graphs and dB graphs on Ig sequences, we employed a second simulation, starting with a single IMGT reference antibody sequence denoted by **𝒜**. Note that an antibody (Supplemental Fig. S7) is a ‘Y’ shape protein consist of a variable region and constant region. The variable region is formed by selecting a gene from each of 3 sets *V, D*, and *J* which are brought together by recombination and splicing. The combined variable region itself can be divided into a framework (FR) which is relatively constant, and three hypervariable complementarity determining regions (CDRs; Supplemental Fig S7) [50]. In the simulation, **𝒜** was created by joining known V, D, and J regions (IGHV1-18*01, IGHD1-1*01, IGHJ1*01; [51]; Supplemental Fig. S8). We generated a collection *D* of decoy sequences in which each nucleotide was chosen uniformly at random, except for the insertion (at a random position) of a single sub-string of **𝒜**. The insertions were of varying lengths ranging from 20 to 26. The antibody reference **𝒜** and decoy gene sequence collection *D* were used as a template to simulate reads, using the tool *wgsim* (https://github.com/lh3/wgsim), with sequence error rate set at *ε* = 1%. A dB graph and an SdB graph was built using these reads to measure the false positive and false negative results from these graphs.

Let *G* = (*V, E*) denote the dB or SdB graph, depending on context, while the graph *G*_*A*_ = (*V*_*𝒜*_, *E*_*𝒜*_) is constructed solely using **𝒜**. In the ideal scenario, *G* and *G*_*𝒜*_ should be identical. Therefore (Supplementary Methods), the false negative rate (denoted by **ℱ**) can be estimated using

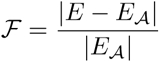

The false positive rate was measured indirectly, using divergence (Supplementary Methods):

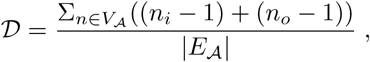

where *n*_*i*_ is the in-degree and *n*_*o*_ is the out-degree of node *n ∈ V*_*𝒜*_. The divergence provides a measure of false connections for the antibody sequence **𝒜**.

Note that false positive edges can also arise due to sequencing errors. However, most dB construction corrects for such errors by choosing an appropriate threshold for coverage, along with other methods [47, 48, 34]. However, the appropriate threshold is different for each value of coverage. We chose a principled method for choosing coverage to remove false positive edges due to sequencing errors for both dB, and SdB. Subsequent to coverage filtering, the false negative rate and divergence was measured as a function of increased coverage (Fig. S9).

Fig. S9 shows an explicit tradeoff between false negatives, and divergence (false positives) for dB graph methods. At any specific fixed coverage parameter(e.g. 10×), the false negative rate of the dB graph increases with increasing values of *k*, even as divergence decreases, making it difficult to simultaneously improve both metrics. In contrast, SdB graphs show consistently lower divergence and false negatives for all coverage values.

### Read filtering

Before we construct the split de Bruijn graph, we need to collect the reads that encode Ig gene transcripts (See Experimental Procedures – ‘Read filter’). We tested the quality of read filtering by a partial alignment of filtered reads to the reference antibody sequences. A virtual antibody reference was set to represent the variable regions of all antibodies. We adjusted the gap between this virtual antibody and each individual antibody using the IMGT antibody reference with gap. The matching *k*-mer was used to anchor the alignment, and the extent of the alignment was determined simply by the length of read on each side of the *k*-mer. The anchored position of the read was transferred to virtual antibody position and used to estimate the overall coverage. We counted the number of reads passing through each unique position of the virtual antibody. Supplemental Fig. S10 describes a coverage due to the partial alignment of all filtered reads, and shows that the reads are filtered without apparent bias except at the very end of the sequence.

### MS-MS based discovery of antibody peptides

We used four mass spectrometry data-sets. To test the algorithms, a data-set of spectra acquired from a purified polyclonal antibody mixture (*antibody purified*) was used [37]. To test for antibody peptides in tumor samples, we used a collection of MS/MS spectra from 90 distinct colorectal tumor samples from the CPTAC project [35] (*colorectal tumor*). As negative control, we used spectra acquired from 30 normal colon biopsies [35] (*colorectal normal*). As a second control, we used spectra from colon cancer cell-lines LIM1215, LIM1899, and LIM2405 (denoted as *colon cell-lines*). [36].

SdB and dB graphs were designed and implemented, utilizing 162.7GB RNA-seq reads of 90 individuals downloaded from The Cancer Genome Atlas (TCGA) repository[52]. The two approaches resulted in a 69.3MB and 107.8MB FASTA-formatted amino-acid database. A multi-stage search (See Experimental Procedures –‘Multi-stage search’) using known proteins and SdB graphs (respectively, known proteins and dB graphs) was conducted to identify peptide spectrum matches (PSMs). A summary of the results of those searches are presented in Table 1. The list of identified spectra and other details are presented in Supplemental Table 1 – ‘Link to the list of PSM and spectrum image’.

Note that the antibody-purified data-set presents an interesting challenge as the SdB graph was constructed from RNA of completely different individuals. Even so, our search identified 16, 404 antibody PSMs (3, 167 peptides) out of 116, 018 total spectra (PSM identification rate 14%). Fig. 1 (a) shows that the identified peptides cover the entire space of the antibody. Table 1 also allows for a comparison of the SdB graph and dB graph databases, as both use the same read set as their input. At identical FDR cut-off (1%), SdB graphs identify 3.3× as many PSMs as the dB for the colorectal tumor, and 1.7× as many PSMs for antibody purified data-set, consistent with simulation results. On the other hand, the number of PSMs identified in the colorectal normal, and cell-line colorectal samples are similar, validating the proposition that SdB graphs can filter out erroneous PSMs at the same rate as dB graphs. Therefore, SdB graphs reduce both false positives and false negatives in the real data, identifying more true PSMs without increasing false PSMs.

In the sample matched colorectal tumor spectra, 54, 909 PSMs (1, 940 peptides) were identified. We asked if these large numbers of antibody peptides originated from tumor infiltrating lymphocytes, or from other sources. For example, these immunoglobulin identifications could simply correspond to floating antibodies from blood contamination, or they could be mis-identified (modified) peptides. In the first case, we would expect to see similar numbers of antibody peptides in colorectal tumor and colorectal normal data set. In the second case, we would expect to see similar numbers of antibody peptides in colorectal cancer, and colon cell-lines (Fig. 2(a)).

We normalized PSM counts to the number of PSMs in the Known DB before comparing across samples (Fig. 2(b)). The normalized PSM count in the colorectal normal data-set was only 4.69% of the colorectal tumor counts (*p*-value *<* 0.0001; See Experimental Procedures –‘Statistical test for antibody enrichment’). The normalized PSM count in colon cell-lines was 0 suggesting that TILs are the source of the antibody peptides observed in the colorectal tumor.

The assembly of RNA-seq reads is a well established research area of genomics [43, 47, 48]. However, genome assembly tools are designed to be general, and may not do a good job of assembling Ig genes. As the reconstruction of Ig genes from the RNA-seq reads was a key part of our pipeline, we asked if the use of the RNA assembly tools could provide better results. To test this, a popular transcriptome assembly tool, rnaSPAdes[49], was used to assemble RNA-seq reads from one colorectal tumor sample. We searched the MS/MS data from the same sample against databases constructed using rnaSPAdes and SdB graphs. The number of spectra identified using the SdB graph method was 2, 450, compared to 528 using rnaSPAdes, suggesting that general purpose transcript assembly tools were not suitable for studying the antibody repertoire at the protein level.

### SAAV peptides

We used the Single Amino Acid Variant (SAAV) peptides reported in previous studies [33, 34], but performed an additional filtering step to keep only the SAAV with the highest confidence. Specifically, we removed all peptides where the mutation has a mass difference of one. We also enumerated all reference peptides with common modifications that shared some sequence tag with the mutated peptide and scored the alternatives. If a reference (modified) peptide could better explain the spectrum, the SAAV was removed from consideration. The final list of 677 mutated peptides is presented in Supplemental Table 3, with the annotated mass spectra in the MassIVE online repository (URL in Supplemental Table 1).

### Antibody-antigen correlation

We asked if the antibody peptides discovered in the colorectal tumor dataset could be targeting specific neo-antigens. The neo-antigens are possibly somatically mutated peptides that are recognized by TILs and antibodies. In previous studies, many somatically mutated peptides had been detected in the same colorectal tumor data by our group, and others [32–35]. Supplemental Table 4 shows the occurrence of all mutated (antigen) peptides, and all antibody peptides in each of the 90 samples.

For every antigen-antibody pair in this table, a calculation was made to determine the significance of co-occurrence using the Fisher exact test. Since many pairs were to be tested, a target-decoy approach was used to compute the false discovery rate for significant pairs. The decoy statistics were computed by permuting the occurrence of each peptide in the sample. Fig. 3 shows the distribution of p-values computed from the target and the decoy table. At a nominal *p*-value threshold of 0.00025, we see 163 pairs that exceed this threshold, versus 5 decoy pairs, suggesting a small false discovery rate of *≤* 5%.

**Figure 3:**
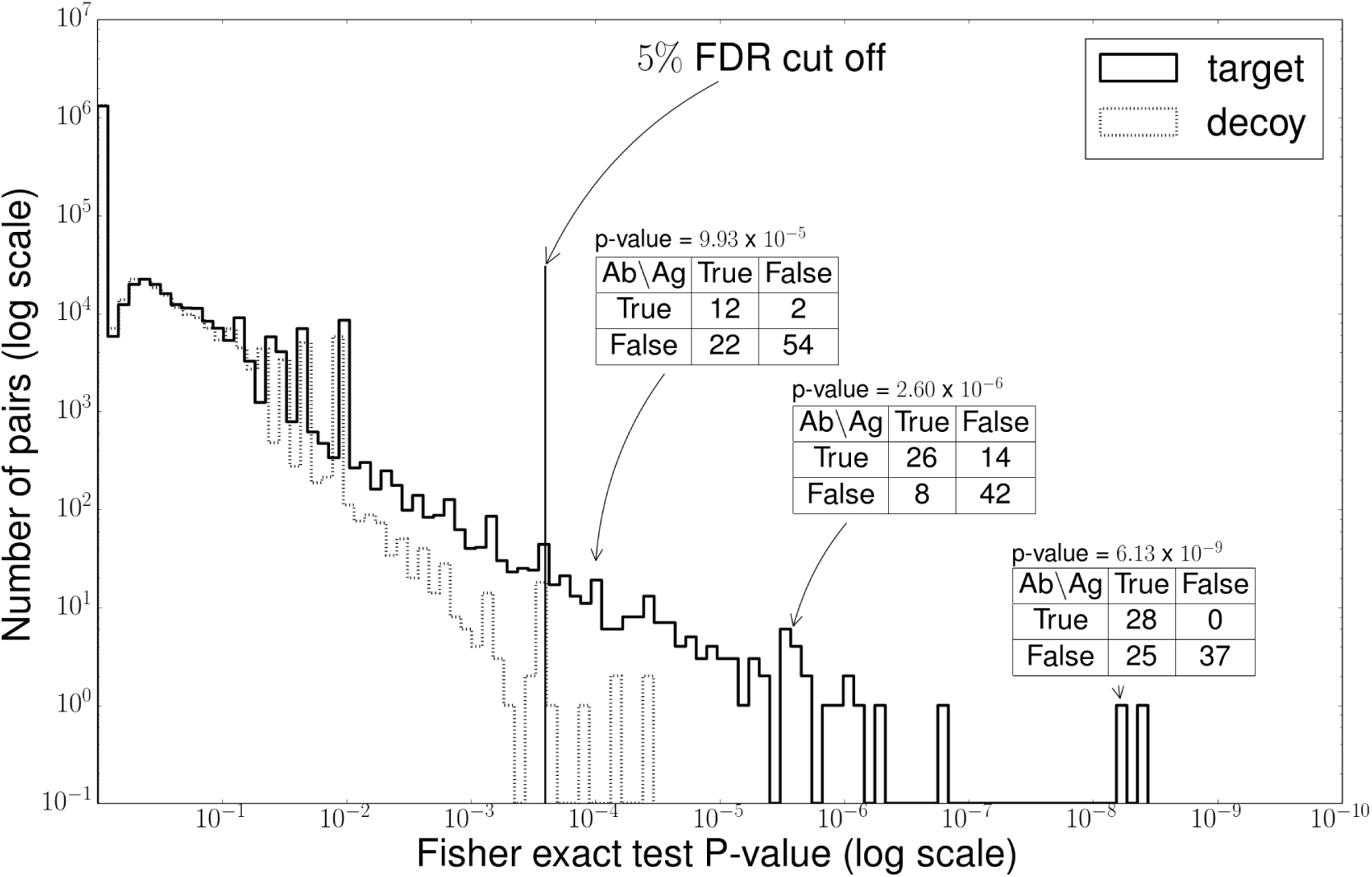
Peptide correlation test. We tested the correlation between the antibody peptides and mutated peptides. For every pair of peptides, we counted the number of samples co-occurring with these peptides and then we applied Fisher exact test to calculate the p-value. For example, the peptide pairs of *N T LY LQM DSLR* (antibody) and *AAQAQGQSCEY SLM V GY QCGQV F* (*Q→R*) (SAAV peptide) cooccurred in 26 samples, and there was a co-absence in 42 samples. It was revealed that 68 of the 90 samples shared the co-occurrence of this pair with a p-value of 2.60 × 10^*−6*^. We drew the histogram of p-values of all pairs in Supplemental Table 4. We also drew the histogram of the p-values from the decoy table generated by the random permutation of values. A 5% FDR threshold was applied to collect the high correlated pairs.

One example of these co-occurring pairs is the the antibody peptide NTLYLQMDSLR, and SAAV peptide AAQAQGQSCEYSLMVGYQCGQVF(Q→R). Note that the antibody peptide NTLYLQMDSLR belongs to variable region of IGHV3-64D*06 and the mutated peptide AAQAQGQSCEYSLMVGYQCGQVF(Q→R) belongs to the gene FBLN1 reported to be down-regulated in colorectal cancer cells [53]. Among 90 samples, both peptides are expressed in 26 samples, and neither is found in 42 samples, giving a Fisher exact test *p*-value of 2.59 × 10^*−6*^. Supplemental Fig. S11 shows examples of peptide spectrum matches of these peptides. These co-occuring antibody, neo-antigen pairs are indicative of an immune response to somatic mutations in tumor genomes. Note that the co-occurence does not indicate co-occurence only between the specific antibody peptide and the SAAV, but rather between the antibody carrying NTLYLQMDSLR and the mutated version of FBLN1 product. In fact we see another antibody peptide LSCAASGFSFR in the FR1/CDR1 region that also co-occurs with AAQAQGQSCEYSLMVGYQCGQVF(Q→R) (*p*-value:9.93 × 10^*−5*^). Therefore, we did not filter antibody peptides by their location (CDR/FR).

### Correlation between antibody expression and survival status

The antibody peptide repertoire might provide a snapshot of the immune response to cancer. We anticipated that the patients with higher immune response could have a different clinical outcomes than those with lower immune response because of the role of TILs in mediating response to cancer[54–56].

We first measured the immune response of an individual as the fraction of identified peptides that came from the antibody repertoire, and identified a subset of individuals as high-responders and low-responders (See Experimental Procedures –‘Measuring immune response’, and Fig. 2(c)). We used the days-to-death values to get the Kaplan-Meier survival estimator for the two groups. Next, we used a log-rank test to compute a *p*-value for the difference between the two curves. The *p*-value was 0.75, indicating that we could not reject the Null hypothesis (Supplemental Fig. S13).

We also considered the possibility that some, but not all peptides mediate a positive clinical outcome. Further, these peptides would be expressed in multiple individuals with similar outcomes. To test this hypothesis, we designed a method that takes any group of peptides, and clusters samples based on co-expression, but without knowledge of the clinical outcome in the individuals (See Experimental Procedures –‘Survival rate comparison’). For a given collection of peptides, we tested the null hypothesis that there is no correlation between sample grouping and the clinical outcome.

We computed an empirical null distribution by choosing random subsets of individuals, and performing the log-rank test against clinical outcome. Supplemental Fig. S5 shows that the test statistic under null hypothesis closely follows the theoretical *χ*^*2*^ distribution.

In contrast, when we tested sample grouping using the correlated antibody, SAAV peptide pairs (See Experimental Procedures –‘Antibody and SAAV peptides correlation test’), we observed a significant differential response with *p*-value: 0.032 (Fig.4 (a)). We also tested this method using two other groups of peptides. When we used all antibody peptides we also obtained a differential response with *p*-value 0.040. (Fig.4 (b)). However, testing with all mutated peptides, we did not observe significant differential response, obtaining a *p*-value of 0.522 (Fig.4 (c)). The small number of samples implies that our study is not fully powered and the results need to be replicated in larger cohorts. Nevertheless, they do show that antibody expression could be correlated with the clinical outcomes.

**Figure 4:**
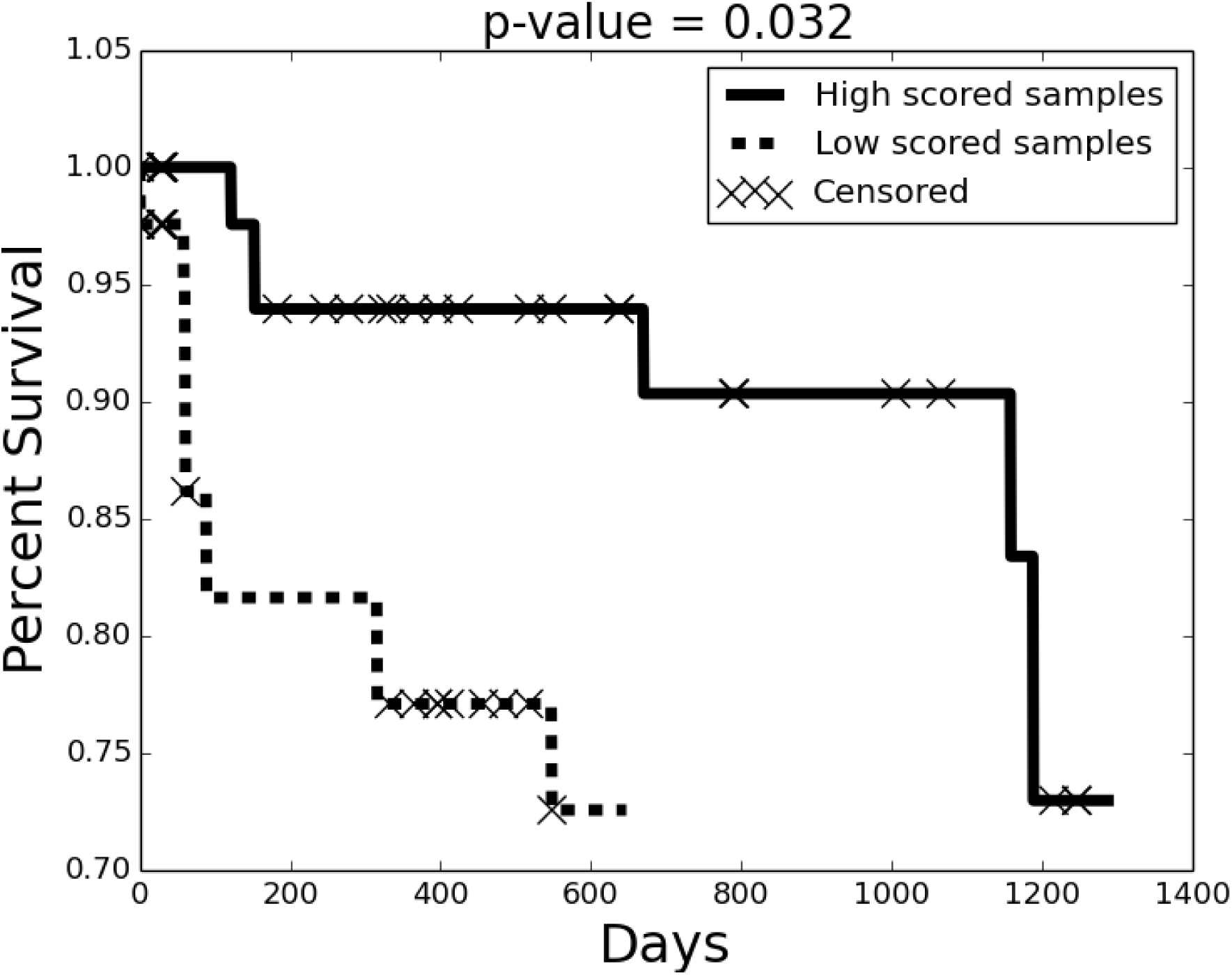
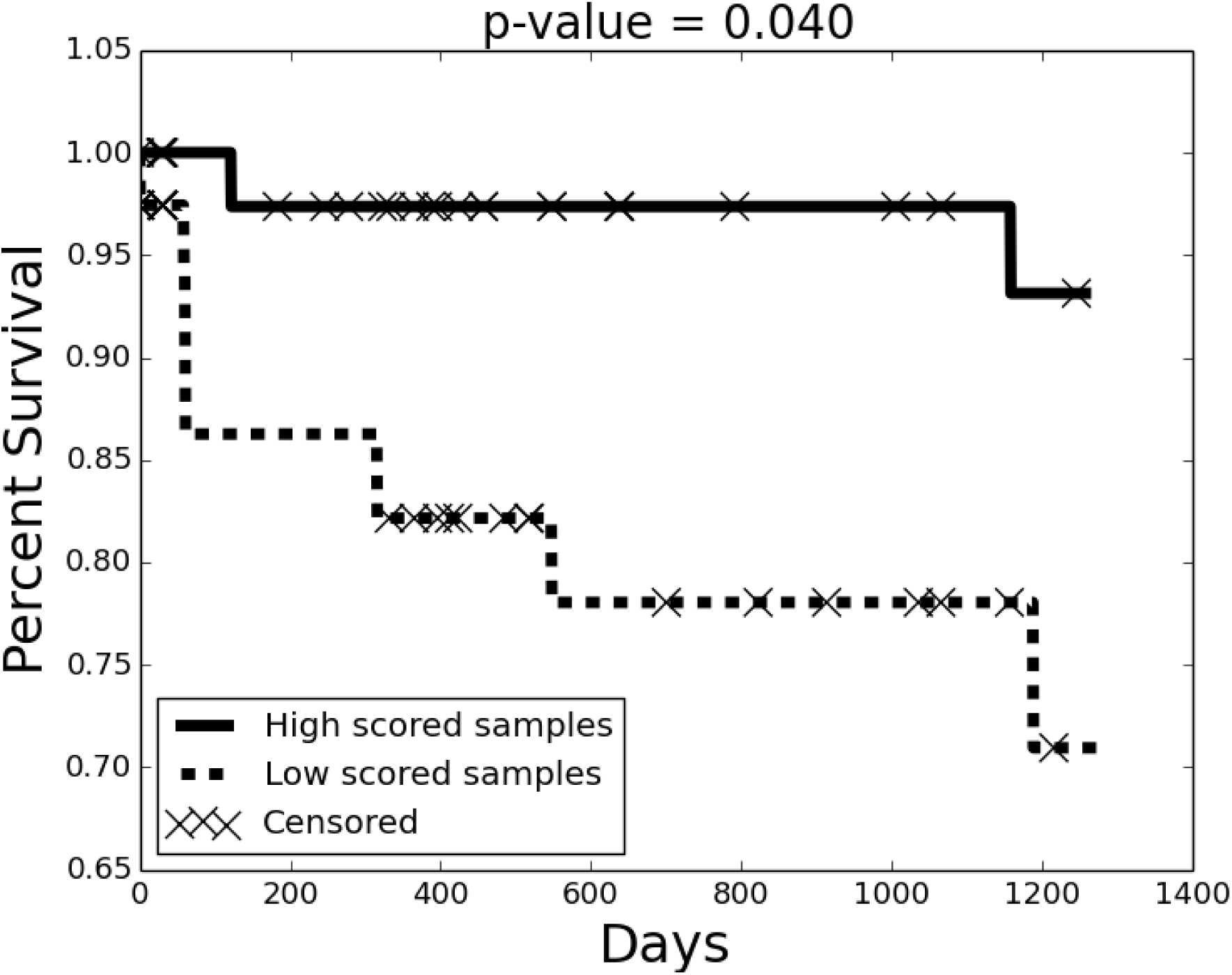
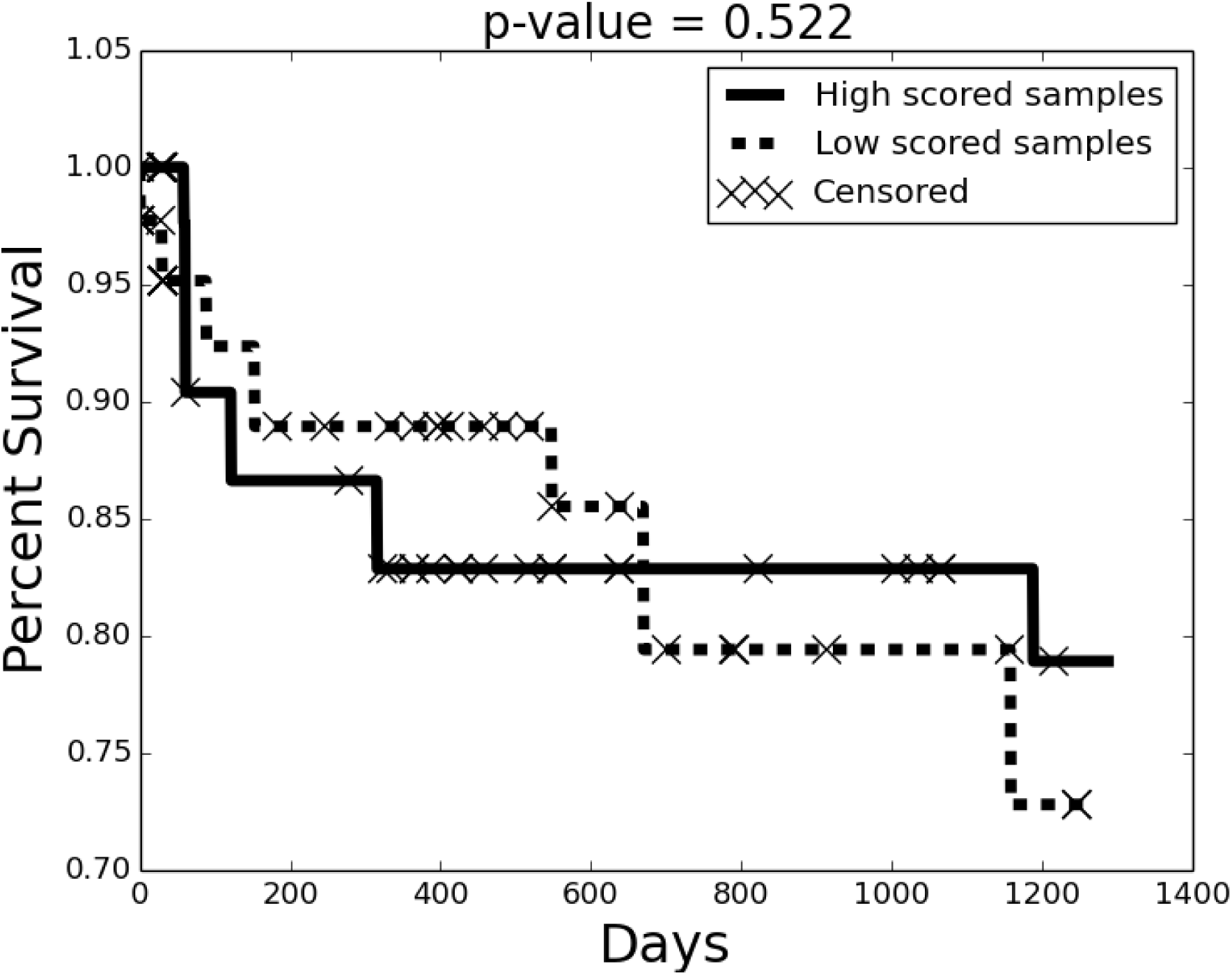
Kaplan-Meier survival estimator. For any subset of peptides, we bi-partioned peptides based on co-expression in samples. Next, we scored each sample based on the homogeneity of peptides from a single partition in that sample (Methods). The highest and lowest scoring samples (one-third each) were grouped, and were tested to determine the clinical outcome. The Kaplan-Meier survival estimator and log-rank test were applied to test the difference of the clinical outcome of two groups. When testing with co-occurring mutated peptide/antibody peptide pairs, we observed a significant correlation with survival (Plot (a): *p*value = 0.032). In contrast, the correlation was reduced when testing with only antibody peptides (Plot (b): *p*-value = 0.040), and there was no-correlation when testing with mutated peptides. (Plot (c): *p*-value = 0.522).

## 7 Discussion and future study

Understanding the immune response to cancer is key to cancer immunotherapy. Current approaches use serum or plasma samples and specifically focus on isolating differentiated B-cells for analyzing antibodies. However, the serum antibody repertoire may contain a larger pool of antibody sequences, not just the ones responding to tumor neo-antigens. In this paper, we mined spectra acquired from isolated (colorectal) tumor cells, and identified a large number of antibody peptides. Our results suggest that infiltrating lymphocytes in the tumors generate antibodies in response to the tumor. They also suggest that somatic coding mutations in the tumor genome act as neoantigens triggering antibody generation. We observed recurrence of antibody and mutated peptide sequences that cannot be explained as chance events, and showed a positive association between clinical outcome (survival time), and the antibody response. Together, the results underscore the need for systematic analysis of the tumor antibody repertoire.

The identification of antibody peptides using tandem mass spectrometry is technically challenging. In the ideal case, the spectra should be searched against transcript data from differentiated B-cells from the same individual. However, that data may not always be available. Moreover, it is not known if circulating B-cells have the same antibody repertoire as the tumor infiltrating lymphocytes. In this paper, we used RNA-seq data, not from isolated B-cells, but from the same tissue that the proteome was extracted. Nevertheless, we managed to get significant coverage of antibody peptides. We identified a large number of peptides even when we used MS data from unmatched samples. Future research will focus on the differences between different sequencing approaches, such as IG-seq, and RNA-seq.

The hyper-variability of antibody sequences makes it challenging to construct databases that can be searched with MS spectra. We proposed a new structure, called the SdB graph, and showed improved performance in compressing and creating MS-searchable databases relative the dB graphs. The SdB graphs are later converted into Fasta formatted databases that can be used for search with any tool. The software for developing SdB graph should be generally applicable for any hypervariable region, and is available for download. These techniques described here can be further improved and those will be the focus of future research.

We found that the SdB graph database generated from RNA-seq of TCGA tumor samples was also helpful in identifying antibodies from completely different samples. This raises the possibility that multiple RNA-seq samples from a specific tumor type could be used as a universal database, reducing the need for matched RNA and protein samples for decoding the immune repertoire. This will be explored in future work. At the end, we also hope that our preliminary results spurs a further investigation of the clinical outcome based on immune system response, and the development of diagnostic tools and therapies that can emerge from an analysis of the tumor immune repertoire.

## 8 Acknowledgement

Dr. Vineet Bafna, Dr. Pavel A Pevzner, and Seong Won Cha were supported by a grant from the NIH (P41RR24851, 1R01GM114362). Dr. Seungjin Na were supported by a grant from the NIH (5P41GM103484-07). We’d like to thank Dr. Alan Carter Covell for his assistance with proofreading the manuscript.

